# Bacterial community response to species overrepresentation or omission is strongly influenced by life in spatially structured habitats

**DOI:** 10.1101/2021.12.01.470875

**Authors:** Hannah Kleyer, Robin Tecon, Dani Or

## Abstract

Variations in type and strength of interspecific interactions in natural bacterial communities (e.g., synergistic to inhibitory) affect species composition and community functioning. The extent of interspecific interactions is often modulated by environmental factors that constrain diffusion pathways and cell mobility and limit community spatial arrangement. We studied how spatially structured habitats affect interspecific interactions and influence the resulting bacterial community composition. We used a bacterial community made of 11 well-characterized species that grew in porous habitats (comprised of glass beads) under controlled hydration conditions or in liquid habitats. We manipulated the initial community composition by overrepresenting or removing selected members, and observed community composition over time. Life in porous media reduced the number and strength of interspecific interactions compared to mixed liquid culture, likely due to spatial niche partitioning in porous habitats. The community converged to similar species composition irrespective of the initial species mix, however, the dominant bacterial species was markedly different between liquid culture and structured porous habitats. Moreover, differences in water saturation levels of the porous medium affected community assembly highlighting the need to account for habitat structure and physical conditions to better understand and interpret assembly of bacterial communities. We point at the modulation of bacterial interactions due to spatial structuring as a potential mechanism for promoting community stability and species coexistence, as observed in various natural environments such as soil or human gut.

**Importance:** Bacteria live as complex multispecies communities essential for healthy and functioning ecosystems ranging from soil to the human gut. The bacterial species that form these communities can have positive or negative impact on each other, promoting or inhibiting each other’s growth. Yet, the factors controlling the balance of such interactions in nature, and how these influence the community, are not fully understood. Here, we show that bacterial interactions are modified by life in spatially structured bacterial habitats. These conditions exert important control over the resulting bacterial community regardless of initial species composition. The study demonstrates limitations of inferences from bacterial communities grown in liquid culture relative to behaviour in structured natural habitats such as soil.

## Introduction

Advances in high-throughput sequencing uncovered vast diversity of microbiomes present in all ecosystems: from soils (Bahram et al 2018, Delgado-Baquerizo et al 2018) and oceans (Ibarbalz et al 2019), to plants and animals (Bai et al 2015), to humans (Costea et al 2018) and urban habitats (Afshinnekoo et al 2015). The important roles that microorganisms play in ecosystem functioning and health have prompted interest in better understanding microbial assembly, stability and activity, towards improved prediction and control of community formation and function (Lawson et al 2019, Widder et al 2016). Resolving the mechanisms that support coexistence of diverse species within a shared environment is particularly challenging, as are the balance of interactions within multispecies assemblages (Little et al 2008) and how such interactions are influenced by habitat characteristics.

Simple experimental ecosystems yield new insights through systematic hypotheses testing and controlled evaluation of theoretical concepts (Cairns et al 2018, Carlström et al 2019, Chodkowski and Shade 2017, Friedman et al 2017, Meroz et al. 2021, Kehe et al 2019, Voges et al 2019). Such approaches have confirmed several essential drivers of community assembly and species interactions, notably the nature and availability of nutrient and carbon resources (Enke et al 2019, Fu et al. 2020, Goldford et al 2018), population density(Abreu et al 2019), rates of migration (Gokhale et al 2018), and how environmental toxicity affects functionality(Piccardi et al 2019). The role of microbial habitat spatial structure has received comparatively less attention, despite well-established understanding in ecological theory that spatially structured environments influence biodiversity, interactions and species coexistence (Nadell et al 2016, Tilman 1994). A few experimental and modelling studies have shown that microhabitat spatial structure could stabilize simple assemblages of 2-3 bacterial species that would not coexist in mixed cultures due to competitive exclusion (Borer et al 2018, Kim et al 2008, Lowery and Ursell 2019, Wang and Or 2013). Therefore, spatial habitat heterogeneity affects bacterial community dynamics in ways irreproducible in homogeneous (liquid) habitats. Here, we hypothesize that spatially structured habitats, such as found in soil and other porous media, dominate the nature of bacterial interactions in space and thus exert a significant influence on bacterial community dynamics. More specifically, porous domains that are partially water-saturated constrain nutrient fluxes and interspecies competition thus providing specific life conditions different from well-mixed habitats (Ebrahimi and Or 2015, Tecon et al 2018, Wang and Or 2013). To test this hypothesis, we have used a simplified bacterial community of 11 species (Kleyer et al 2019) in replicate microcosms supplied with the same nutrient resources but with variation in spatial structure and hydration state (Fig. 1). The porous habitats comprised of glass beads were set at controlled hydration states from wet to relatively dry, while liquid media without beads served as contrasting unstructured habitats. The synthetic community comprises 11 members from phyla commonly found in soil (*Proteobacteria, Actinobacteria* and *Firmicutes*). Functional traits with relevance for the soil environment include decomposition of organic matter, nutrient cycling, and maintenance of soil fertility. In particular, the community members play an important role in nitrogen fixation (*Paenibacillus sabinae, Pseudomonas stutzeri*, *Rhizobium etli*, *Xanthobacter autotrophicus*), are involved in bioremediation of different components like phenols (*Arthrobacter chlorophenolicus* (Westerberg et al 2000)), polychlorinated biphenyl (PCB) (*Burkholderia xenovorans* (Liang et al 2014)), polycyclic aromatic hydrocarbons (PAHs) (*Pseudomonas stutzeri* (Singh et al 2017)) and halogenated hydrocarbons (*Xanthobacter autotrophicus*). We added *Escherichia coli* as non-soilborne species to the community as *E. coli* often ends up in the environment as faecal contaminant (Brennan et al 2010). All members can grow aerobically and some are able to enter dormancy by forming spores (*Bacillus subtilis, Paenibacillus sabinae,* and *Streptomyces violaceoruber* (Brenneret al (2005)). Thus, the selected synthetic community covers a wide phylogenetic range and combines bacterial species with different life strategies. To uncover interactions within this community, we manipulated the initial composition of the inoculum by systematically over representing one species at a time. Our working hypothesis was that spatially structured habitats would select for a bacterial community composition that differs from liquid culture habitats, moreover, we predicted that the resulting community composition in structured habitats would be marginally affected by the initial inoculum with overrepresented species. To elucidate the types of interactions (bacterial competition or facilitation), we selectively removed species from the inoculum (one at a time). Our high-throughput system and the modular approach using defined species and habitats is well suited for disentangling abiotic factors and identifying interspecies interactions, the two key components for scrutinizing mechanisms of bacterial community assembly.

**Figure 1.**
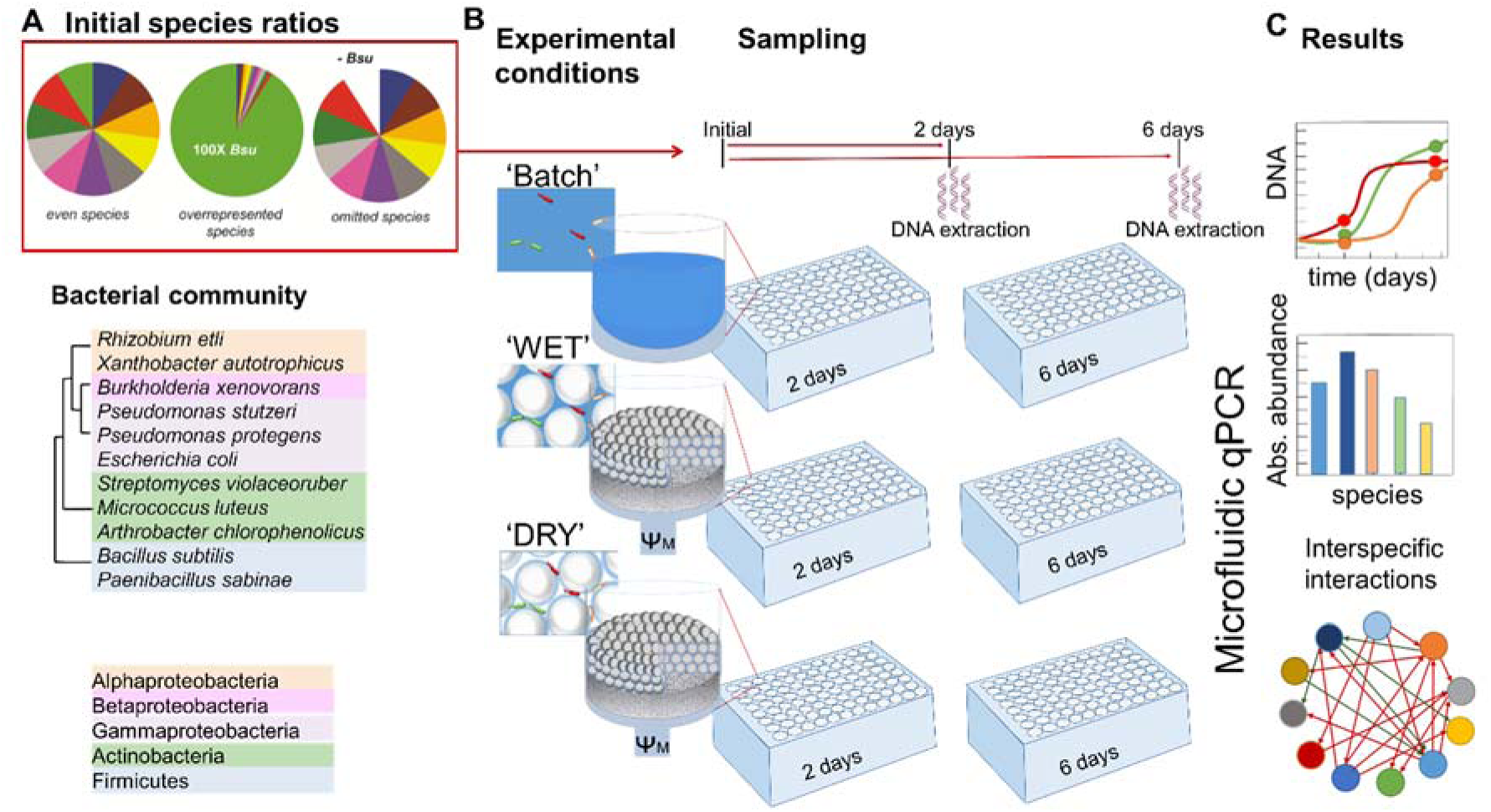
Synthetic ecology as experimental approach. (A) Synthetic bacterial community of 11 phylogenetically diverse species. We manipulated the initial species ratio in the community as follows. Species were mixed in even proportions, in uneven proportions with one species being a 100-fold overrepresented, and in even proportions but with one species removed. An example with *B. subtilis* is shown. (B) Microcosms were set up in multi-well plates containing only liquid medium (‘batch’), or liquid medium and glass beads (diameter ≈100 µm) to provide a spatial structure to the habitat, with prescribed hydration status in the microcosms permitting us to maintain relatively ‘wet’ or ‘dry’ conditions in the structured habitats. Even and uneven species mixes were used as inocula. Two multi-well plates per treatment were prepared, and sacrificed after 2 and 6 days of incubation. Beads and bacteria are shown for illustration purposes and their relative size is not on scale. (C) Microfluidic-based qPCR was used to resolve bacterial community composition at the species level to obtain relative and absolute abundance data.

## Results

### Convergence of bacterial community composition in structured habitats

We manipulated the initial composition of the synthetic bacterial community inoculum (Supplementary Table S2) by targeted overrepresentation and omission of selected members and followed the development of community composition growing in an unstructured batch habitat and structured microcosms kept under ‘wet’ and ‘dry’ conditions (Fig. 1). The absolute abundance for each community member, estimated by the number of species ‘genome equivalents’ (≈number of bacterial cells, see *Materials and Methods*) in each microcosm shows an increase of the total community under all conditions from inoculation to 2 days and further at 6 days of incubation (Fig. 2). The resulting species abundance varied by several orders of magnitude from virtually undetected to 8×10^8^ cells per microcosm (representing 3.2 ×10^9^ cells per g glass beads) for the most abundant species (Fig. 3). Community composition in all microcosms were dominated by gammaproteobacteria (*Pseudomonas protegens*, *Pseudomonas stutzeri*, *Escherichia coli*), irrespective of time or initial species ratio (Fig. 3). Remarkably, the absolute and relative abundance of these three dominant species was consistent across replicates and determined primarily by the presence or absence of microcosm spatial structure and by the hydration status of structured habitats. While *P. protegens* and *E. coli* dominated liquid batch microcosms, *P. stutzeri* was the most abundant species in microcosms containing glass beads, with this trend increasing over time (Fig. 3). The marked changes in relative abundance of dominant species between structured and non-structured microcosms seem to be controlled by the growth of *P. stutzeri*, whose abundance was poor in batch (10^4^-10^5^) but highest of all species in glass bead microcosms both under wet or dry conditions (≈10^8^, an increase of 3-4 orders of magnitude). Spearman’s rank correlations between the Bray-Curtis similarities of microbial community composition in all samples confirmed *P. stutzeri* as main driver for the observed patterns (73 %). Importantly, the divergence of community compositional patterns observed in structured and non-structured microcosms was not driven by the total size of the bacterial community, which was highest in wet structured habitats (up to 10^9^ cells per microcosm) and similarly lower in liquid and dry structured habitats (Figs. 2A and 3). Principal coordinate analysis (PCoA) of Bray-Curtis dissimilarities (Fig. 2B) shows a distinct cluster of bacterial communities from liquid habitats separated from communities in structured microcosms. The structured habitats communities were grouped based on hydration condition (‘wet’ and a ‘dry’ clusters). Additionally, the different sampling times (2 days and 6 days) formed delimited sub-clusters within their respective hydration conditions (Fig. 2 B). High similarity of community structure within the same habitat at 2 and 6 days are further confirmed by ANOSIM pairwise comparison of each habitat and time point (Fig. 2 C). Moreover, ANOSIM statistics also underscore strong differences in community structure between batch culture incubation and communities from structured microcosm habitats.

**Figure 2.**
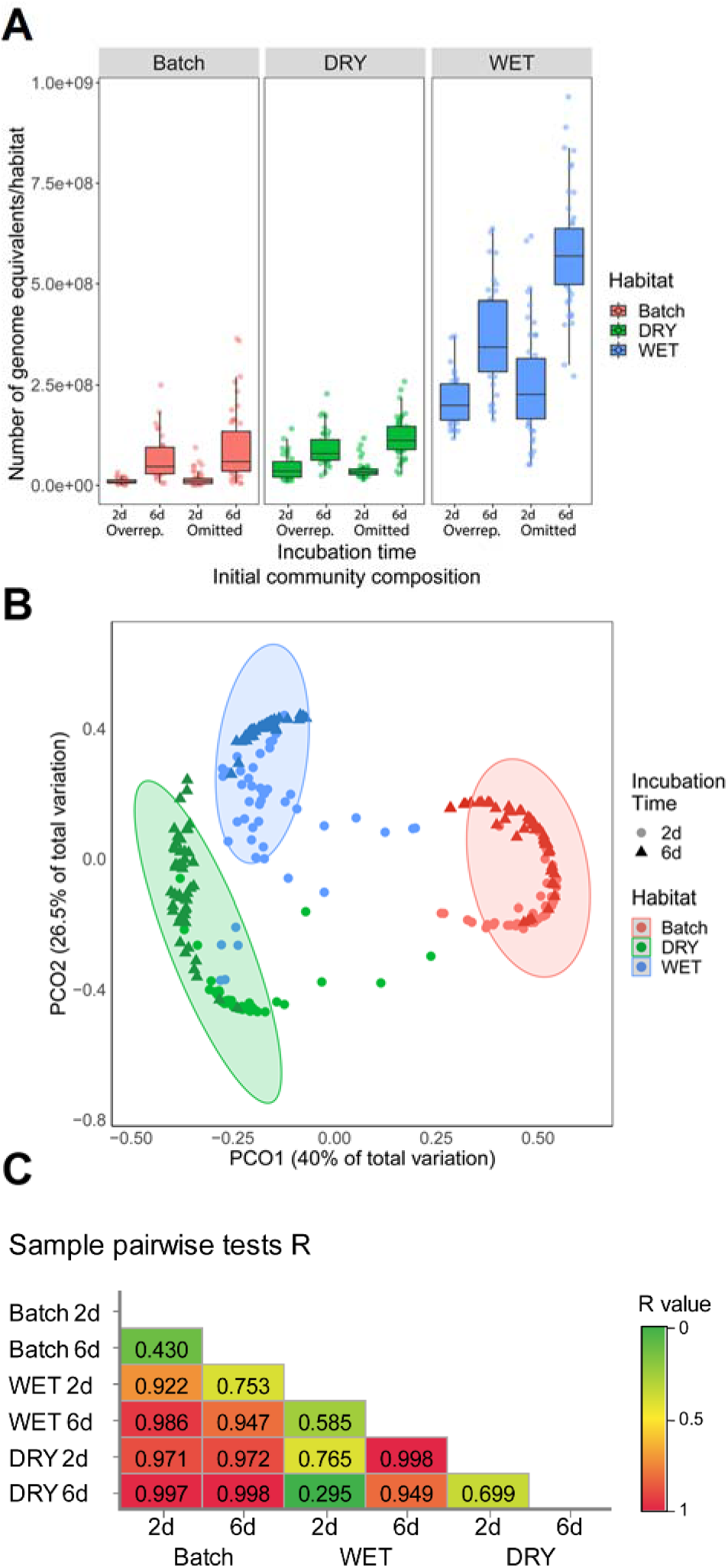
(A) Total bacterial community size as function of treatment and time. Boxplots show the estimated sum of all species in the community, expressed as number of genome equivalents per microcosm. Total abundances were systematically higher in wet structured microcosms compared to dry structured and batch microcosms, and after 6 days compared to 2 days. Omitting or 100x overrepresentation (Overrep.) of a species did not appear to affect the total counts. (B) PCoA plot based on Bray-Curtis dissimilarity calculated for microcosms inoculated with an even mix of species or mixes of species containing an overrepresented member. Clustering indicates the largest separation between communities incubated in unstructured batch microcosms and in glass-bead structured microcosms under WET and DRY conditions, ellipses enclose 95% of the data points for each condition. Further separation is observed for the time of sampling (2 days and 6 days) forming partially overlapping sub-clusters. (C) Pairwise comparisons of dissimilarity between treatments were calculated with ANOSIM. R values close to 1.0 (red) suggest high dissimilarity between treatment groups. Communities in batch habitats strongly differ from communities growing in structured microcosms under DRY and WET conditions. Moreover, differences in community structure were observed between communities from structured microcosms kept under DRY and WET conditions. Community structure within the same treatment at the early and late time point show intermediate R values indicating some overlap.

**Figure 3.**
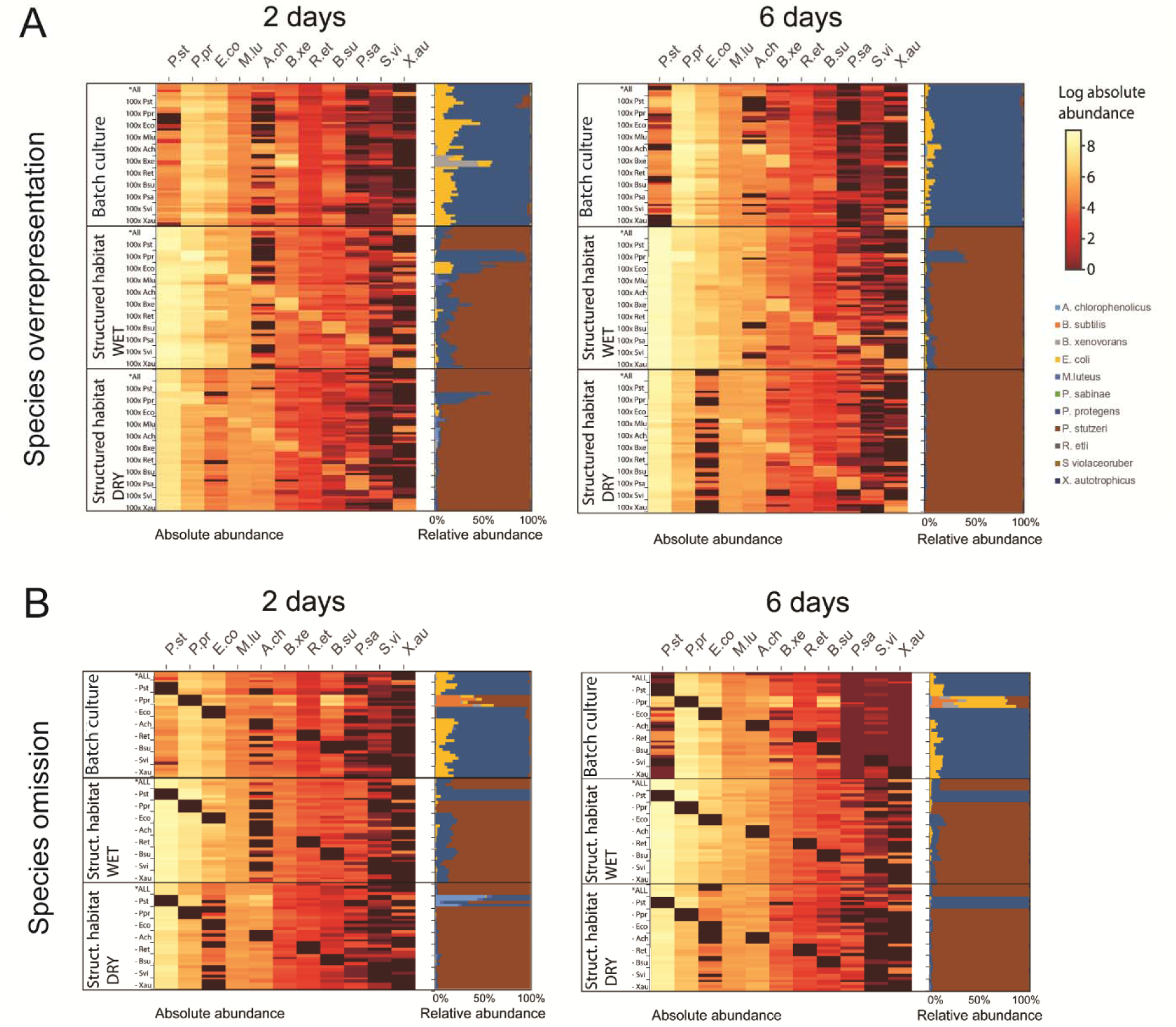
Variations in bacterial community composition in microcosms. The synthetic bacterial community was grown in multi-well plates containing only liquid medium (liquid habitat) or liquid medium mixed with glass beads (spatially structured habitats), the latter with prescribed hydration providing constant wet or relatively dry conditions. Microcosms were sacrificed after 2 or 6 days of incubation for total DNA extraction. Heatmaps show the absolute abundances (log-transformed) of each species (columns) based on the absolute number of genome equivalents measured in each microcosm by qPCR. Absolute counts were used to calculate relative abundance values, shown alongside heatmaps. (A) We investigated the effects of initial species ratios by manipulating the initial composition ratios of the inoculum (100-fold increase of one species at a time compared to the others, 100X…), while a community with even proportions of all species served as a control (*All). The different overrepresented inocula are shown as rows, each with four replicate microcosms per mix. (B) We selected 8 species to be removed from the initial inoculum, thus allowing us to test for causal effects on the growth of the remaining species. The communities are shown as rows, with four replicate microcosms per mix.

### Manipulation of initial species ratio has limited effects on community convergence

The overrepresentation of individual species in the initial inoculum (Fig. 1) did not affect the overall dominance of gammaproteobacteria in unstructured and structured habitats (Fig. 3A). While overrepresentation resulted in significant changes in community composition, those changes were transient and waned or disappeared after 6 days of incubation (most notably in the case of *P. stutzeri*, *P. protegens*, *E. coli* and *Burkholderia xenovorans*). After 6 days, the relative abundance patterns of the dominant species were virtually unaffected by a 100-fold increase in initial abundance of each species compared to the others, with the exception of *P. protegens* that persisted at high abundance at 2 and 6 days when overrepresented in structured habitats (Fig. 3A). Similarly, in most cases the patterns of dominant species persisted despite omission of selected species in the initial inoculum. Nevertheless, marked changes occurred when one of the members of the dominant trio of gammaproteobacteria was removed (Fig. 3B). Specifically, the removal of *P. stutzeri* in structured habitats led to increased *P. protegens* growth and relative abundance, which produced a pattern similar to that of liquid habitats. *Arthrobacter chlorophenolius* was also stimulated by the removal of *P. stutzeri*, but only in dry structured habitats, and the effect was no longer visible after 6 days. *P. protegens*, once removed from liquid habitats, was replaced by a diversity of other species (*P. stutzeri*, *E. coli, A. chlorophenolicus, B. xenovorans, Bacillus subtilis*) with high reproducibility among replicate microcosms (Fig. 3B).

### Signatures of initial species overrepresentation persists in structured habitats

We systematically tested the consequences of a 100-fold increase of one species over the others, one species at a time, in the mix inoculum. Although the relative abundance patterns of dominant species persisted, as discussed above, imprints of this initial manipulation of community composition were detected in the absolute abundance counts. These compositional signatures were prominent in structured habitats (wet and dry), exhibiting a ‘staircase’ pattern of absolute abundance heatmap (Fig. 3A). Two days after inoculation, this imprint was visible for all 11 species in the community in structured habitats, whereas in liquid habitats its persistence was limited to five species (*P. stutzeri, A. chlorophenolicus, B. xenovorans, B*. *subtilis* and *Xanthobacter autotrophicus,* Fig. 3A). Moreover, this characteristic pattern was confirmed by the analysis of significant effects of one species initial overrepresentation over the same species final abundance (Fig. 4A and Supplementary Fig. S2) with significant increase for all 11 overrepresented community members (p < 0.05) in structured wet habitats (Fig. S2).

**Figure 4.**
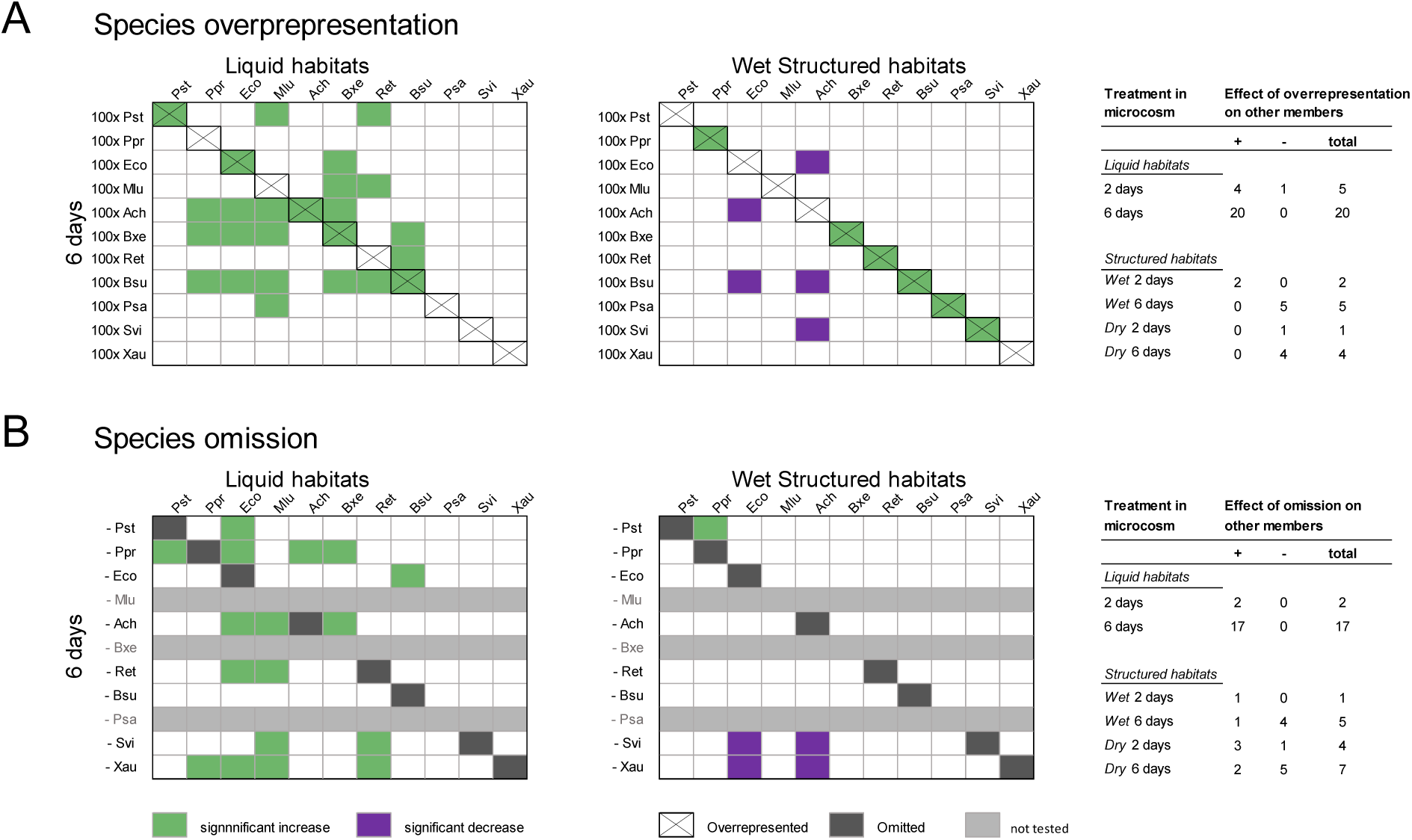
Positive and negative effects of species overrepresentation and omission on other species. Microcosms were inoculated with initial community mixes containing a 100-fold overrepresented species (A) or one omitted species (B). For simplicity we present results from liquid and wet structured habitats after 6 days (full results are shown in Supplementary Fig. S2). Green and magenta colors indicate that the absolute abundance of a given species was respectively significantly higher or lower as a response to overrepresentation or omission of another species in the mix compared to its abundance in control microcosms (inoculated with an even mix of all species, either in liquid or in porous habitats). Variations (increase or decrease) in absolute abundances were deemed significant when >5-fold with a p-value <0.05 (using a two-tailed t-test on 4 replicate microcosms). Tables show the detected effects for all treatments and time points, excluding the effects of a species on itself. A significant increase (+) or decrease (-) of one species as a response to overrepresentation or omission of another is interpreted here as an interaction between the two species, which could be competitive or facilitative depending on the context.

### Species overrepresentation or removal from inoculum uncovers interspecific interactions

We further analyzed the absolute counts data to identify causal relationships between the overrepresentation or omission of a given species and the absolute increase or decrease of another bacterial species abundance (measured as number of genome equivalents), which we define here respectively as positive and negative effects. We compared species abundances in microcosms inoculated with uneven initial species mixtures with abundances in microcosms inoculated with an even mix of species serving as a control (Fig. 1B), for each treatment and time point. Selected results presented in Fig. 4 (and full results in Supplementary Fig. S2) show the detected changes in the abundance of one species with respect to the control (with a >5-fold change threshold contingent to a p-value<0.05). The total number of detected positive and negative effects in liquid habitats was higher than in structured habitats in both cases of species overrepresentation and omission (Fig. 4 and Supplementary Fig. S2). While the number of detected positive effects increased with time in liquid habitats, it decreased in structured habitats (Fig. 4). Negative effects (i.e., decrease in another species’ abundance) were observed for one single case in liquid habitats at day 2 (decrease of *B. subtilis* in response to overrepresentation of *P. protegens*), but were frequent in structured habitats, especially after 6 days (Fig. 4 and Supplementary Fig. S2). Results obtained from species omission suggested more and stronger competitive interactions in liquid than in structured habitats (Fig. 4B and Supplementary Fig. S2). After 6 days in liquid, the removal of *P. protegens* had permitted significant growth increase in five other species: *A. chlorophenolicus*, *B. subtilis*, *B. xenovorans*, *E. coli* and *P*. *stutzeri*, but this was not the case in structured habitats (Fig. 4B and Supplementary Fig. S2).

We identified a number of reciprocal responses to overrepresentation and omission, that is, instances where a species both increased in absence and decreased in overrepresentation of another species (or vice versa): *E. coli* in response to *S. violaceoruber* and *X. autotrophicus* (wet structured); *A. chlorophenolicus* in response to *P. stutzeri* (dry structured); *B. subtilis* in response to *P. protegens* (liquid).

## Discussion

The study highlights dynamic adjustments in species composition of a well-defined bacterial community (Fig. 1) grown in different habitats with two distinctly different outcomes: (1) a community dominated by *P. protegens* for liquid habitat; or (2) a community dominated by *P. stutzeri* in porous and structured habitat (Fig. 3). Under all conditions gammaproteobacteria dominated the microcosms, which suggested a faster or more efficient use of the nutrients available in the growth medium. However, the difference between (1) and (2) could not be attributed to nutrient resources (invariant across microcosms), nor to the total growth of the community (Fig. 2A). We propose that, to account for this discrepancy, the nature of the habitat is the primary determining factor (i.e., the presence or absence of physical structure with microscopic spatial arrangement of habitats) (Fig. 3A). Generally, the results supported the hypothesis that spatially structured habitats exert strong influence on growth, interactions and assembly of bacterial communities. Additionally, the nature of the habitat and interactions is not only determined by the solid phase (glass beads), but by the hydration state (aqueous phase) as well. The organization of the aqueous phase near water saturation (‘wet’) or under relatively drier conditions (‘dry’) had a significant influence on species abundance. With ‘wet’ porous habitats giving rise to bacterial community composition similar to liquid habitats, with higher abundances of *P. protegens* and *E. coli* and lower abundance of *A. chlorophenolicus* (Fig. 2A). This result emphasizes the importance of aqueous phase organization on habitat connectivity and community composition, as also supported by previous experimental and modelling studies (Borer et al 2019, Kim and Or 2017, Kleyer et al 2019).

The use of a synthetic bacterial community enabled systematic manipulation of initial species composition in microcosms (i.e., with 100-fold relative increase or removal of individual species, Fig. 1). Our results showed that irrespective of initial bacterial species ratio, the community composition drifted from an initially even abundance to similar patterns dominated by *Pseudomonas* species. Even after 6 days, the initial differences in species ratio did not alter the relative abundance patterns with respect to the most abundant species in the community (except when the dominant species *P. protegens* or *P. stutzeri* were omitted from the inoculum) (Fig. 2). Remarkably, the imprint of initially overrepresented bacterial species was preserved in communities grown in porous media with structured habitats (Fig. 2A, Fig. 4A, and Supplementary Fig. S2). This suggests that physical structure and aqueous habitats forming between glass beads may either stimulate species growth or reduce interspecific interactions irrespective details of the hydration status. This translates to a stabilizing effect of structured environments that preserves initial community composition for extended periods relative to community composition in liquid culture with potential implications for engineering stable microbial communities (Ben Said et al 2020, Lindemann et al 2016).

In addition to bacterial-habitat interactions, we seek insights on interspecific interactions that shape community composition and the roles of community members and their collective functioning. Selective initial overrepresentation or omission triggered significant changes in different species’ final abundances (Figs. 2 and 4). In a background of similar resource availability, we link these changes to the onset of competitive or facilitative interactions between various species during growth (Bruno et al 2003, Ghoul and Mitri 2016). Competition is expected to dominate interactions in our system, because all species grow independently to high cell densities in the complex liquid medium (a nutritious mix of protein digests with glucose and mannitol as additional carbon sources) (Kleyer et al 2019). This is supported by the drift from an initially even community composition to a highly uneven composition observed in all treatments after 2 days, with gammaproteobacteria out-competing other phyla (Fig. 2). It is further supported by the omission of one of those dominant species resulting in higher abundances of similar competitors, notably the replacement of *P. protegens* by *P. stutzeri* and vice versa (the two *Pseudomonas* species are closely related phylogenetically and share certain traits such as fast growth on rich media) (Fig. 4B). In addition to competing for nutrients and space, the bacterial species in the community might also directly antagonize each other (Little et al 2008). *P. protegens*, in particular, is known to secrete a variety of antimicrobial metabolites (Ramette et al 2011) and possesses a Type VI secretion system that is involved in contact-dependent elimination of competitors (Vacheron et al 2019). Such antagonizing mechanisms could potentially also explain the marked effects of *P. protegens* omission in liquid cultures (Fig. 4B). By contrast, facilitation (e.g., metabolic cross-feeding between species) is less expected considering that the growth medium is complex (thus obscuring trophic dependencies (D’Souza et al 2018)) and non-toxic (Piccardi et al 2019). Overall, our results from species omission experiments supported these expectations on the nature of interactions in our system (Fig. 4B, Supplementary Fig. S2). The use of a synthetic ecology approach was here decisive to clearly disentangle species interactions from habitat filtering (Ghoul and Mitri 2016, Vorholt et al. 2017); the impossibility to do so remains an inherent limitation in network analyses of microbial communities from natural habitats (Röttjers and Faust 2018).

The number of detected changes suggesting species interactions was overall higher and largely positive in liquid compared to structured habitats (Fig. 4 and Supplementary Fig. S2). We found that in response to species overrepresentation in liquid habitats, other phylogenetically distant members exhibited an increase in abundance (e.g., increase of *B. subtilis* or *B. xenovorans* facilitated growth of five community members from four different phyla), which could indicate potential cross-feeding due to metabolic dissimilarities. Somewhat surprising are results from species omission that suggest prevalence of facilitative interactions in structured habitats, while results from species overrepresentation suggested the converse (Fig. 4 and Supplementary Fig. S2). However, we note that overrepresenting and omitting species are not strictly symmetrical manipulations, and in that context we consider species omission to be a more definitive arbiter in determining species interactions (see also for example Gutiérrez and Garrido, 2019).

Importantly, our results demonstrate that porous and structured microhabitats modulate the number, type and strength of interspecies interactions. These effects have been previously suggested for small bacterial assemblages of 2-3 species (Borer et al 2018, Kim et al 2008). As noted above, most natural habitats harbour complex spatial structures that likely mitigate competitive effects. This understanding challenges a widespread (albeit debated) view that competition dominates interactions among microorganisms in nature (Foster and Bell 2012). We surmise that the experimental reliance on liquid cultures to study microbial interaction processes may overemphasize the prevalence of interspecies competition within microbial communities in nature (Foster and Bell 2012, Rivett et al 2016).

The role of spatial structures highlights certain limitations of the widespread use well-mixed liquid media to study bacterial interactions and community assembly. The simplicity and reproducibility offered by such media comes with a bias of the behaviour of such communities in natural and structured environments (i.e., soil or human gut). These differences are rooted in the complex spatial arrangement of solids, liquid and gas phases at the microscale and their impacts on the diffusion, motility and interspecific interactions in most natural environments (Tecon and Or 2017, Vos et al 2013). Consequently, we argue that the physical spatial structures of microbial habitats is a central element necessary for any accurate prediction of bacterial community dynamics and stability, alongside well recognized factors such as the type and diversity of carbon sources (Goldford et al 2018, Zegeye et al 2019).

Arguably, a fundamental goal of microbial ecology is to uncover the ecological mechanisms that permit the coexistence of many diverse species within a shared environment (Tilman 1982). Among the identified mechanisms, niche partitioning is often predicted to play a major role (Saleem et al 2015). Niche partitioning can refer to differential use of resources (‘ecological’ or ‘metabolic’ niche) (Saleem et al 2015) and to segregation in distinct spatial areas (‘spatial’ niche) (Ghoul and Mitri 2016). We conclude from our results that the latter is more likely to be at play in structured microcosms (containing glass beads at various hydration states), hence supporting species coexistence. In general, when considering that many natural habitats are porous media (notably, soil) characterized by physico-chemical gradients and spatial structures, we may contemplate a vast diversity of spatial niches arising that are absent in liquid environments (Bickel and Or 2020, Raynaud and Nunan 2014, Tecon and Or 2017). As noted above, spatial structuring leads to another complementary mechanism promoting bacterial species coexistence: the abatement of strong ecological interactions (positive or negative) between species that perturb the community. This mechanism was notably proposed to account in part for the observed stability of the human gut microbiome (Coyte et al 2015).

Results presented here underscore the crucial role of habitat structure, key for our understanding of processes in natural bacterial communities inhabiting heterogeneous environments where structures define and stabilize composition and dynamics that essentially impact community functioning (e.g. gut- and soil microbiota). Lasting effects from directed manipulation of bacterial community composition are fundamental for communities used as probiotics, in food production or for environmental restoration, remediation and in biocontrol. Linking effects of hydration and the physical spatial structure with dynamics in a defined bacterial community offers a useful tool in sustainable engineering of community composition.

## Materials and Methods

### Bacterial strains, culture conditions and synthetic community assembly

We assembled a synthetic bacterial community comprised of 11 species listed in Fig. 1. Bacterial strains were obtained from the Leibniz-Institute German collection of microorganisms (DSMZ): A6 (DSM12829), 168 (DSM402), LB400 (DSM17367), MG1655 (DSM18039), DSM20030, T27 (DSM17841), CHA0 (DSM19095), CMT.9.A (DSM4166), CFN 42 (DSM11541), A3(2) (DSM40783), 7C (DSM432). All bacterial strains were cultured on 0.1x tryptic soy broth (TSB) agar plates (VWR International, Leuven, Belgium) supplemented with 1% mannitol (TSBM) at 25 °C (mannitol sustains the growth of *R. etli*). For inoculation, 48h-old plates were scraped with a sterile spartel and bacteria were suspended in PBS (Phosphate Buffered Saline) solution with pH 7.4 (Gibco, life technologies Europe, Bleiswijk, Netherlands). Bacterial biomass was measured using optical density at 600 nm (OD_600_). *Rhizobium etli* and *Xanthobacter autotrophicus* secreted copious amount of extracellular polymeric substances (slime) when grown on agar plate, and for that reason these two species were subjected to an additional washing step in PBS in order to remove slime prior to OD_600_ measurement. *Streptomyces violaceoruber* grows as a mycelium, which can bias OD_600_ measurement. Therefore, *S. violaceoruber* was further treated in an ultrasonic bath (Branson Ultrasonics, Danbury, Connecticut, United States) at 40 kHz for 30 seconds in PBS to homogenize the bacterial suspension prior to OD_600_ measurement. All bacterial communities were prepared with the same final size of approx. 5.5x 10^4^ cells (calculated from OD_600_ measurement) with PBS used for dilutions. For the community containing all 11 members, individual species with an OD_600_ of 0.1 were combined in equal proportions and further diluted with PBS to approx. 5 000 cells of each member resulting in a total community size of ∼5.5x 10^4^ cells. For the overrepresentation one member was added with 100x fold higher initial abundance (∼50 000 cells) to the other ten species with ∼500 cells each (total community size 5.5x 10^4^ cells). Inocula missing one species were prepared by combining 10 out of 11 members at approx. 5’500 cells each, resulting in communities of ten members with a total community size of ∼5.5x 10^4^ cells. As culture medium, 240 µl of 0.1x TSBM were added to each microhabitat and microcosms were inoculated with 10 µl of the bacterial community. The same procedure was applied to inoculate liquid cultures in 96-well system kept in parallel to the experimental set-up at constant temperature of 25 °C.

### Bacterial community growth in microcosms

For details on microcosms preparation and incubation, see Supplementary Methods. Briefly, structured microcosms were set-up in multi-well plates with a 0.22 µm filter membrane at the bottom of the well (Merck, Darmstadt, Germany) and containing 250 mg of sterile glass beads (with diameter of 80 to 120 µm) per well. Microcosms were connected via a saturated porous plate to a sterile medium reservoir containing 800 ml of saline solution. The height of the liquid column (the difference in elevation between the microcosms surface and the liquid medium level in the reservoir) prescribed the liquid tension and thus the hydration conditions in the porous plate and the glass-beads microcosms. In parallel, conventional multiwell plates were used for homogeneous liquid microcosms. Each multiwell plate contained 80 parallel microcosms with 20 different initial inocula. An inoculum with the 11 species at equal proportions; 11 with one member 100x overrepresented and 8 with one member missing with four replicate wells per community inoculum. Two liquid habitat multiwell plates and four structured habitat multiwell plates (2 WET and 2 DRY, Fig. 1) were prepared. One plate of each treatment was sacrificed after 2 and 6 days of incubation at 25 °C for total DNA extraction from microcosms.

### DNA extraction and quantification

The microcosms contents were transferred by pipetting to individual 1.2 ml tubes (Brand GmbH + co KG, Wertheim, Germany), immediately frozen in liquid nitrogen and stored at -80 °C. At the end of the experiment all samples were thawed on ice, and DNA was extracted using the DNeasy blood and tissue kit (Qiagen, Hilden, Germany) following the manufacturer’s instructions for increased cell lysis efficiency. DNA concentration was quantified fluorometrically using the Qubit dsDNA HS assay kit (Thermo Fischer Scientific). See Supplementary Methods for details.

### Microfluidic quantitative real-time PCR

Microfluidic quantitative PCR was performed using a Fluidigm 192×24 Dynamic Array (Fluidigm Corporation, San Francisco, CA, USA). Details on the qPCR method using assay chips with integrated fluidic circuits (IFCs) for high throughput is described in previous work (Kleyer et al 2017, Kleyer et al 2019), including a nested-PCR approach where the 16S rRNA gene is preamplified with universal primers to enhance the signal for real-time PCR quantification of individual community members via species-specific PCR primers (Supplementary Table S1). For more details, see Supplementary Methods.

### Data Analysis and Statistics

Real-time quantitative PCR data were analysed with the software provided by the manufacturer (Fluidigm Corporation). We used standard curves obtained from a mixture of pure genomic DNA from each species to quantify the absolute abundance of individual species in the microcosms (Supplementary Fig. S1). This absolute abundance was estimated for a given species as the number of ‘genome equivalents’ per unit mass genomic DNA based on information on each species genome size (for more details see reference (Kleyer et al 2019)). Total species abundance in each microcosm was back-calculated from total community DNA extracted per microcosm (Qubit high-sensitivity assay for dsDNA; Thermo Fisher Scientific) (Fig. 2A). Measurement values <1 genome in DNA template were not reported. A Bray-Curtis dissimilarity matrix was calculated in R (www.r-project.org) with the R package vegan. To show most of the variation we performed a principle coordinate analysis PCoA (R-package vegan) and visualized the result with ggplot2 (Fig. 2B). Analysis of similarity (ANOSIM) was performed with the R-package vegan to obtain statistical R-values for dissimilarity between time points and treatments (Fig. 2C). Interspecific interactions reflected in positive and negative effects of overrepresented- or omitted community members were calculated from absolute abundance data by comparing the abundance of each species per sample to the abundance of the same species in the control microcosms (inoculated with an even mix of all species). Variations (increase or decrease) in absolute abundances were deemed significant and thus represented when >5-fold with a p-value <0.05 (using a two-tailed t-test on 4 replicates).

## Acknowledgments

We thank Aria Minder and Silvia Kobel from the Genetic Diversity Center of ETH Zurich for help with the Fluidigm high throughput qPCR. Financial support for this work came from the Swiss National Science Foundation (project number 31003A_182734). We declare that the research was conducted in the absence of any commercial or financial relationships that could be construed as a potential conflict of interest.

## Author Contributions

H.K., R.T. and D.O. designed the research. H.K. performed the experimental research and analyzed data. H.K., R.T. and D.O. wrote the manuscript.

## Supplementary Information

### Supplementary Methods

#### Preparation and incubation of microcosms

Structured microcosms were set-up in 96-well plates with a flat 0.22 µm filter membrane at the bottom (Merck, Darmstadt, Germany) containing 250 mg of sterile glass beads (with diameter of 80 to 120 µm) per well. Individual 96-well plates were placed on a layer of quartz flour atop a ceramic plate (1 bar, 12 cm diameter, 0.5 cm thick). The quartz flour ensured a tight connection of the plate’s wells with the ceramic -plate. Both the layer of quartz flour and the ceramic plate had been pre-saturated with saline solution (0.9% NaCl) and connected to a medium reservoir containing 800 ml saline solution in a bottle. All parts had been sterilized by autoclaving at 120 °C (saline-solution, medium, glass beads) or alternatively treated at 100 °C for 20 minutes (ceramic disc with heat sensitive glue). The saturated quartz flour and ceramic plate permitted us to maintain a continuous liquid connection between the microcosms and the medium reservoir, and the height of the liquid column served to prescribe hydration conditions in the microcosms. Lowering the reservoir bottle induces drainage of liquid from the microbead habitat by a suction that increases with increasing vertical distance between the bead surface and the liquid level of the medium reservoir. This method allows us to induce unsaturated conditions where the energy level of the water is reduces near particle surface where capillary and adsorptive forces dominate. The matric potential (ᴪ_m_) describes adhesive intermolecular forces between the water and the solid as negative pressure expressed in –kPa with the equation ᴪ_m =ρ_ g h, where ρ is the density of water, g is the acceleration of gravity, and h is the height of the liquid column (1). Microcosms in this experiment were kept at a fixed hydration level mimicking WET and DRY conditions with a matric potential of – 0.5 kPa and -6 kPa that is ~5 cm and ~60 cm head difference between the microcosm surface and the medium level in the reservoir. Matric potential was allowed to establish and equilibrate for 5 h prior to inoculation.

Two liquid habitat set ups (2 day and 6 day sampling point) and four microbead setups (2 days WET + DRY and 6 days WET + DRY) and were prepared. All multiwall plates harbour 80 parallel microcosms for 20 different microbial communities (one at equal proportions, 11 with one member 100x overrepresented and 8 communities missing one members) and four biological replicates per community. Structured habitats containing 250 mg glass-beads were placed on top of the hydration controlled ceramic. Fast wet up of the glass-beads indicate connectivity of the system. As culture medium 240 µl 0.1x TSBM were added to each microhabitat and bacterial communities were inoculated as described above.

#### DNA extraction and quantification

For nucleic acid purification the bacterial communities were recovered from the microcosms including the liquid medium or the microbead-matrix and suspended in 180 µl lysis buffer for nucleic acids purification (ATL buffer), to ensure recovery of filamentous growing species a cut tip with enlarged aperture was used. After transfer to 96-rack with 1.2 ml tubes (Brand GmbH + co KG, Wertheim, Germany) the samples were immediately frozen in liquid nitrogen and stored at -80 °C. After the experiment was completed samples were thawed on ice and DNA was extracted using the procedure described in the DNeasy blood and tissue handbook (Qiagen, Hilden, Germany) and the customized additional protocol for increased cell-lysis efficiency through a bead-beating step initiating the extraction procedure provided by the manufacturer. For cell lysis 250 mg glass beads were added to the liquid culture samples (WET and DRY samples already contained the glass bead-matrix) and samples were homogenized in a TissueLyser for 30 s at 6.5 m s−1. DNA was eluted with 50 µl elutions buffer AE to increase DNA yield a second elution step was performed using the 50 μl eluate from the first elution. DNA concentration was quantified fluorometrically using the Qubit dsDNA HS assay kit (Thermo Fischer Scientific) designed specifically to detect double-stranded DNA with high sensitivity in a 384 well plate assay. For readout, we used the Spark 10M Multimode Microplate Reader (Tecan, Männedorf, Schwitzerland) (Fig. S1).

#### Microfluidic quantitative real-time PCR

DNA concentration was normalized to approx. 4 ng/µl with a pipetting robot in a liquid handling station (Brand GmbH + co KG). For the pre-amplification PCR, with 9.6 µl of the community DNA in a total reaction volume of 20 µl containing 200 nM of forward and reverse primers (Supplementary Table S1) in 1x GoTaq G2 Colorless Master Mix (Promega, Duebendorf, Switzerland). Preamplification was performed on a PCR thermal cycler (Sensoquest, Goettingen, Germany) with the following program: initial denaturation at 95°C for 10 min, followed by 15 cycles of denaturation at 95°C for 30 s, annealing at 59°C for 30 s, elongation at 72°C for 90 s, followed by a final elongation step at 72°C for 10 min. Primers were removed prior to reatl-time quantitative PCR by treating 5 μl of the reaction with 10 U Exonuclease I (Thermo Fisher Scientific) at 37 °C for 15 min followed by a heat inactivation step of the enzyme at 85 °C for 15 min. The remaining PCR reaction was amplified for another 15 cycles, and the product was further visualized on an agarose gel to control for the presence of a single amplicon with size of ~1.5 kb.

The exonuclease-treated PCR products were diluted 5 times in DNA suspension buffer (10 mM Tris, pH 8.0, 0.1 mM EDTA) and 2 µl diluted PCR product were added to 3 µl sample mix containing 1x qPCR Mix (HOT FIREPol EvaGreen qPCR Mix Plus (ROX), Solis Biodyne, Tartu, Estonia) and 1x DNA Binding Dye (Fluidigm Corporation), with a final volume of 5 µl. For the assay mix 1x Assay Loading Reagent (Fluidigm, PN 85000736) was combined with 100 μM of forward and reverse primer and nuclease free water was added to a final volume of 4 µl (3 µl per inlet plus 1 µl overage). Supplementary Table S1 contains species-specific primer pairs used in the assay. All primers were previously designed and tested for the characterization of our representative soil bacterial community.

After mixing 3 μl of the assay and of the sample mix were added to the respective inlet of the 192.24 Dynamic Array (Fluidigm) and the chip was loaded and run on the Biomark system using the same real-time quantitative PCR protocol as described previously (2) In total samples from 480 bacterial communities (4 biological replicates) plus 20 community DNA samples used for inoculation were assayed on 3 IFC-chips each designed for 192 samples to be automatically combined with a duplicate set of 12 primer-pairs (Gene Expression 192.24 IFC, Fluidigm corp.) together with two negative controls and an eight point standard calibration in duplicate per chip.

## Supplementary Figures

**Figure S1.**
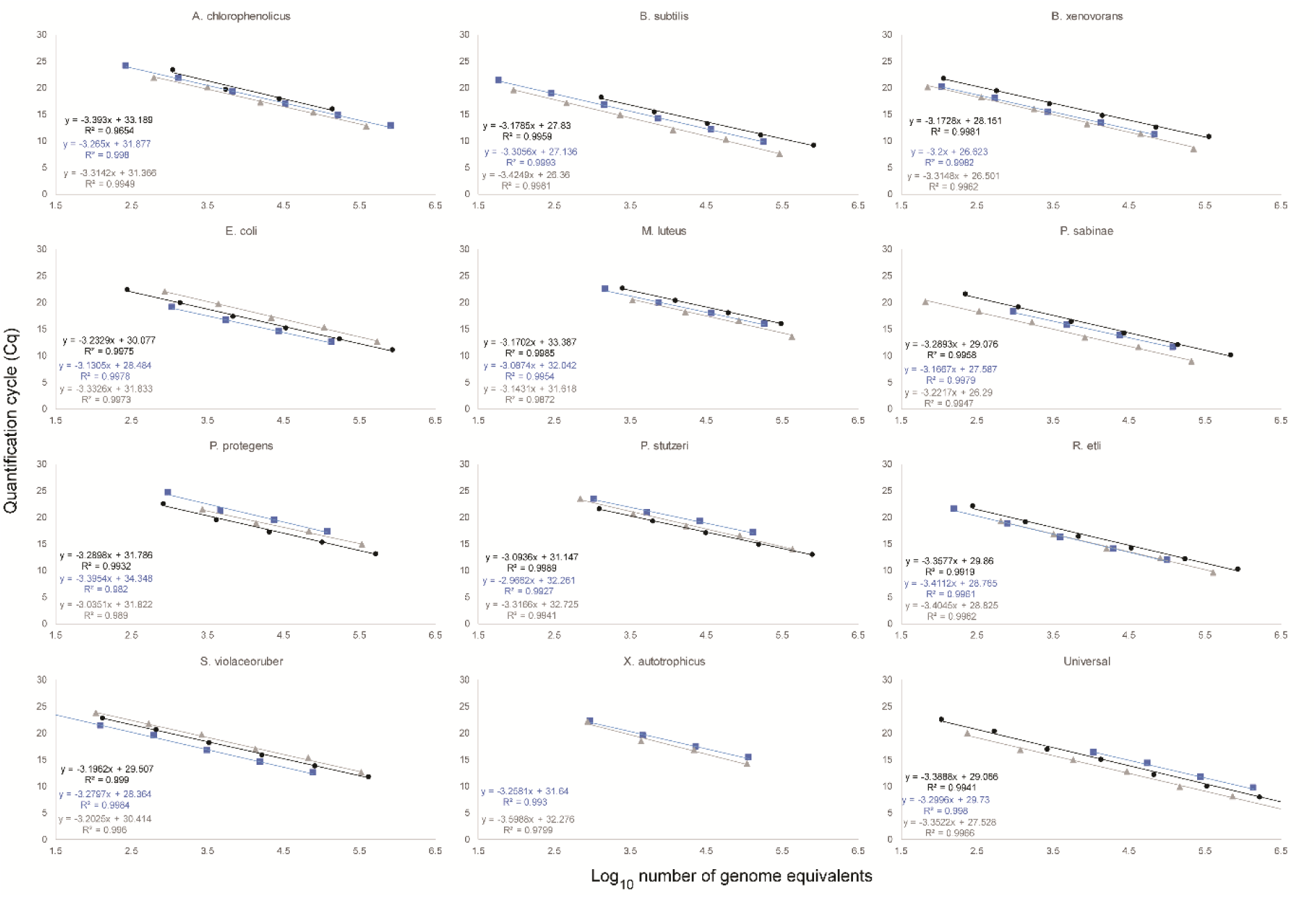
Standard Calibration Curves. Standard calibration curves to calculate abundance of each member of the bacterial community from qPCR data. Purified genomic DNA with known concentration from eleven individual species (Bacterial strains detailed in the Material and Methods section) were combined in equal proportions to prepare fivefold serial dilutions and run in parallel reaction on a Biomark GeneExpression 48.48 IFC (Fluidigm) with each individual species-specific primer pair or with a universal primer pair (Table S1). Based on genome size and known concentration of genomic DNA per species in the stock solution the number of genome equivalent copies was calculated and plotted against the cycle threshold for each species-specific qPCR assay. Data evaluation was performed with the Fluidigm software from four technical replicates per dilution. Standard calibration was calculated for three independent qPCR chips, displayed in black, blue and grey. Equations of fitted linear regression lines and R2 values are shown, calculated from average Cq values. For data evaluation the average values from all three calibrations were used.

**Figure S2.**
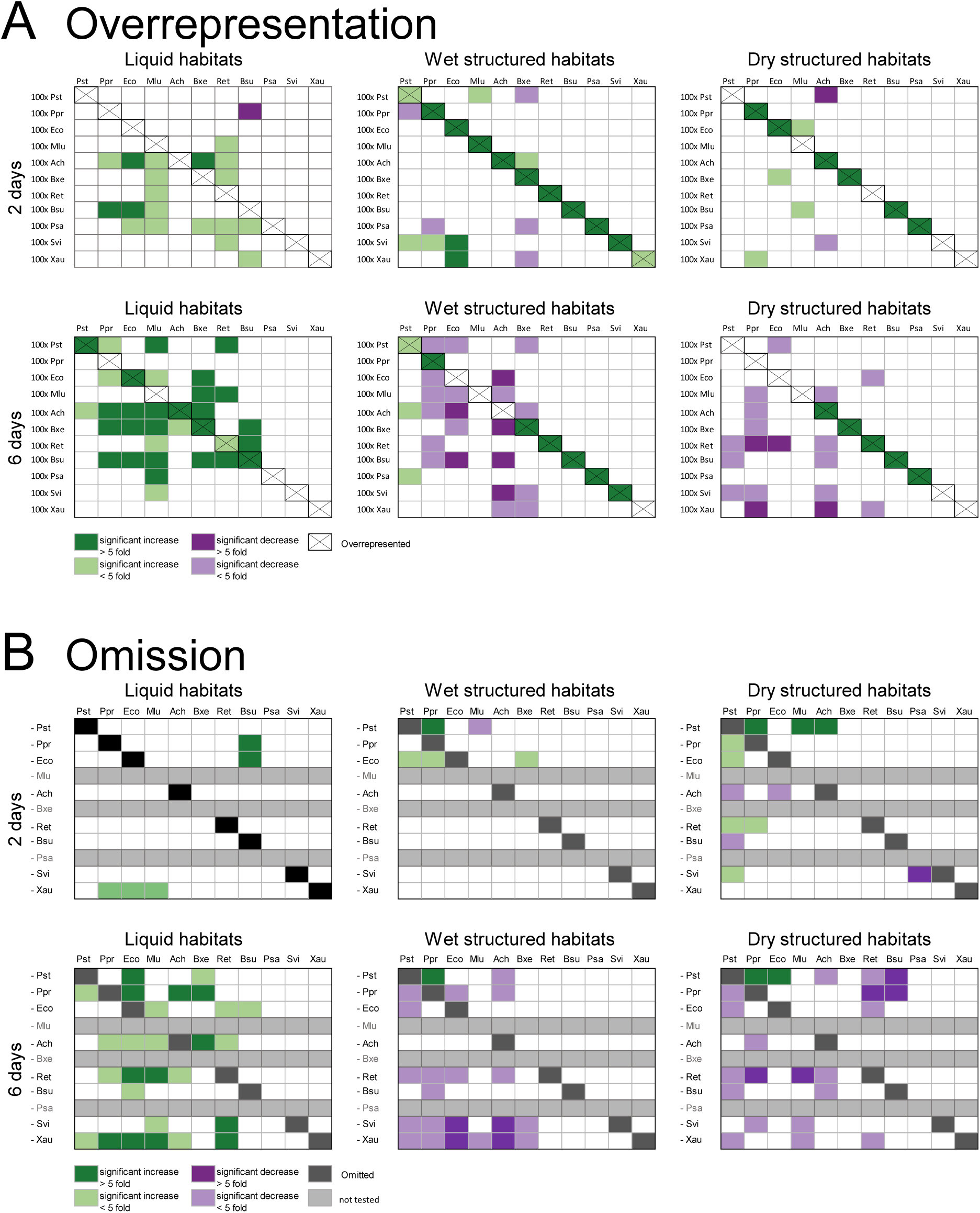
Positive and negative interactions between individual species. (A) Microcosms were inoculated with initial community mixes containing one overrepresented species. (B) Microcosms were inoculated with community mixes where one species has been omitted. This was done for all species except *B. xenovorans*, *M. luteus* and *P. sabinae*. Green and magenta colors indicate that the absolute abundance of a given species was respectively significantly higher or lower compared to the control microcosms (inoculated with an even mix of all species). Variations (increase or decrease) in absolute abundances were deemed significant with a p-value <0.05 (using a two-tailed t-test on 4 replicates) for light colours and significant with an in-decrease of >5-fold dark colours.

## Supplementary Tables

**Table S1.**
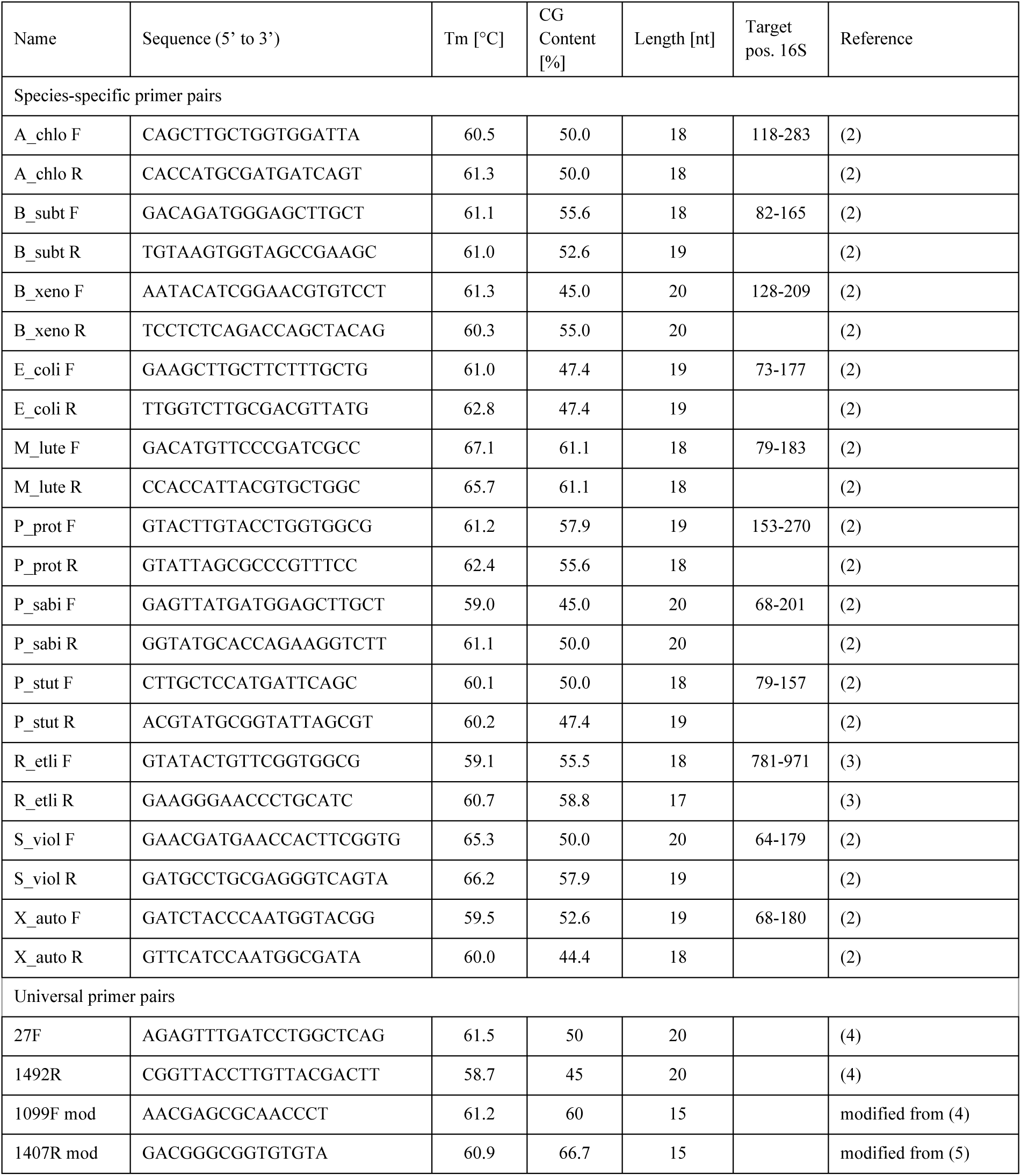
Primers used in real-time PCR.

**Table S2.** Species used in study are well characterized at the genomic and phenotypic level and span a wide diversity of bacterial phyla. Selected species differ in physiology, but all grow aerobically and can be cultivated under standard laboratory conditions.

| Species | Phylogeny/Class | Description |
| --- | --- | --- |
| *Arthrobacter chlorophenolicus* (Ach) | Actinobacteria | Soil, rods/cocci, motile, non-spore-forming, involved in bioremediation of phenols such as 4-chlorophenol (4-CP), 4-nitrophenol (4-NP) and phenol [6] |
| *Bacillus subtilis* (Bsu) | Firmicutes, | Soil, rods, motile, spore-forming, of industrial relevance due to production of hyaluronic acids, specifically proteases [7, 8] |
| *Burkholderia xenovorans* (Bxe) | Betaproteobacteria, | Rhizosphere, rods, motile, non-spore-forming, play important role in bioremediation due to ability to degrade polychlorinated biphenyl (PCB) [9] |
| *Escherichia coli* (Eco) | Gammaproteobacteria | Usually found in intestine of warm-blooded organisms, but often detected as faecal contaminant in environmental samples [10, 11], rods, motile, non-spore-forming |
| *Micrococcus luteus* (Mlu) | Actinobacteria | Soil cocci, non-motile, non-spore-forming, involve in bioremediation: ability to degrade toxic organic pollutants and tolerance to metals [12] |
| *Paenibacillus sabinae* (Psa) | Firmicutes | Rhizosphere, rods, motile, spore-forming, have ability to fix nitrogen [13] |
| *Pseudomonas protegens* (Ppr) | Gammaproteobacteria | Rhizosphere, rods, motile, non-spore-forming, have biocontrol properties: plant-protecting bacteria, produces the antimicrobial compounds pyoluteorin and 2,4-diacetylphloroglucinol (DAPG) which are active against various plant pathogens [14] |
| *Pseudomonas stutzeri* (Pst) | Gammaproteobacteria | Rhizosphere, rods, motile, non-spore-forming Nitrogen fixation, denitrification, bioremediation due to ability to degrade polycyclic aromatic hydrocarbons (PAHs) [15] |
| *Rhizobium etli* (Ret) | Alphaproteobacteria | Rhizosphere, rods, motile, non-spore-forming, form symbiotic relationship with legumes performing nitrogen fixation [16] |
| *Streptomyces violaceoruber* (Svi) | Actinobacteria | Soil, filaments, non-motile, spore-forming, model organism for gram-positive soil bacteria of high G+C content that undergoes a complex life cycle of mycelial growth and spore formation and produces a variety of antibiotics and other drugs during the differentiation process [17] |
| *Xanthobacter autotrophicus* (Xau) | Alphaproteobacteria | soil, rods, non-motile, non-spore-forming Nitrogen fixation, involved in bioremediation via degradation of 1,2-dichloroethane, methanol and propane [18] |

